# Depression-like behavior following mild traumatic brain injury in adolescent rats: a role for limbic neuropeptides

**DOI:** 10.1101/2025.10.16.682872

**Authors:** Cydney R. Martin, Laura L. Giacometti, Jessica R. Barson, Ramesh Raghupathi

## Abstract

Mild traumatic brain injury (mTBI) is common among adolescents because of their participation in contact sports and mTBI is more likely to lead to depression-related behaviors in girls than boys. Various neuropeptides, often within the limbic system, have been implicated in the regulation of depression-related behaviors. To identify potential neuropeptide involvement in behavioral effects of mTBI, this study used adolescent-age male and female rats to compare the effects of single and repetitive mTBI on depression-like behavior and limbic neuropeptide expression. Female but not male rats displayed increased immobility in the forced swim test compared to sham-injured rats at 5-weeks (chronic), but not 2-weeks (acute) following closed-head injury, and this effect was estrous cycle-dependent following single but not repetitive injury. In the nucleus accumbens (NAc), a major limbic nucleus, mRNA expression of corticotropin-releasing factor (CRF) and dynorphin (DYN) was decreased after single injury only in female rats, particularly during the estrus phase, while expression of enkephalin (ENK) was decreased after repetitive injury. In the paraventricular nucleus of the thalamus (PVT), another limbic nucleus, mRNA expression of pituitary adenylate cyclase-activating polypeptide (PACAP) was increased in repetitively injured female but not male rats at 2- and 5-weeks but was unchanged in single injured female rats compared to sham-injured rats. These data suggest that depression-like behavior emerges in the chronic phase following adolescent mTBI only in females, being estrous cycle-dependent following single-injury. Moreover, reduced ENK in the NAc and elevated PACAP in the PVT may contribute to depression-like behavior in females following repetitive mTBI.

## Introduction

One to two million traumatic brain injuries (TBIs) diagnosed in the emergency department annually are classified as mild.^1,2^ Mild TBI (mTBI) can result in problems with memory and concentration, anxiety and depression, and headaches, which can persist in a subset of patients.^3^ Athletes who play contact sports are particularly vulnerable to sustaining repeated mTBI, exacerbating the associated injury deficits.^4–6^ Female athletes typically exhibit different symptoms and take longer to recover, with menstrual cycle stage contributing to this phenomenon.^5,7–9^ The vulnerability of female athletes is further magnified in adolescents, as this population may take longer to recover than other age groups and may not exhibit symptoms until years after injury.^10–12^ Thus, adolescent females with a history of mTBI are more than three times as likely to receive a depression diagnosis, with depression beginning 6 or more months following injury.^13,14^

Few studies have investigated depression-like behavior after mTBI in both male and female adolescent rodents.^15–18^ At 2 weeks after a single or repetitive lateral mTBI, adolescent female rats displayed differences in time spent immobile in the forced swim test (FST),^17^ while 4 days after repetitive injury, adolescent mice showed no change in behavior in the FST.^18^ In contrast, we have demonstrated that 6 weeks following either a single or repeated mTBI during adolescence, injured female but not male Sprague-Dawley rats display greater immobility in the FST, an effect that is dependent on estrus phase only after the single injury.^15,16^ The specific mechanistic basis for these injury-induced sex differences remains to be understood.^15–18^

Neuropeptides expressed in the limbic system can regulate both cognitive and emotional behaviors^19–21^ and several have been evaluated in the acute period following TBI.^22–25^ Within the nucleus accumbens (NAc), dynorphin (DYN) and enkephalin (ENK) are major neuropeptides found in medium spiny neurons^26^ while corticotropin-releasing factor (CRF) is found more sparsely,^27^ but these neuropeptides regulate depression-like and other affective behaviors, often in response to stress.^28–33^ Immunoreactivity for DYN is increased in the brain immediately following repetitive blast mTBI in adult mice, and pre-treatment with an antagonist for the DYN receptor reduces anxiety-like behaviors.^25^ Immunoreactivity for ENK is increased in the pituitary of adult male rats following a severe TBI.^34^ Within the paraventricular nucleus of the thalamus (PVT) which sends dense projections to the NAc, pituitary adenylate cyclase-activating polypeptide (PACAP) is highly expressed^35–37^, which has been implicated in depression-like behaviors in rats and mice.^38,39^ Expression of PACAP is increased in the perifocal cortical region and dentate gyrus within a few to 72 hours following moderate TBI,^40^ and intracerebroventricular injection of PACAP-38 prior to TBI reduces motor and cognitive deficits and neuronal apoptosis.^22^

Expression of CRF receptor mRNA is increased in multiple extrahypothalamic limbic regions following blast-induced mTBI in adult mice, along with an increase in anxiety-like behavior,^24^ and intracerebroventricular injection of a CRF receptor 1 antagonist in adult male mice exposed to a lateral mTBI reduces anxiety-like behaviors.^23^ Female rodents have shown differences compared to males and across estrous phase in behavior and brain circuit activity to central infusion of CRF,^41,42^ in their sensitivity to a DYN receptor agonist in the NAc in ethanol drinking behavior^43^ and they have higher overall expression of PACAP within the PVT.^35^ Thus, neuropeptides in the limbic system may underly sex-related differences in behavioral sequelae following mTBI.

The objective of the present study was to determine the effect of mTBI sustained in adolescence on depression-like behavior in adulthood and of the associated expression of neuropeptides within the NAc and PVT. Importantly, this study sought to investigate sex differences as well as the effect of single and repetitive mTBI in these effects.

## Materials and Methods

### Subjects

Fifty-nine male and 107 female adolescent Long-Evans rats (35 days old, Charles River Laboratories, Wilmington, MA) were used in this study and housed in a 12-hour light/dark cycle with *ad libitum* access to food and water. Animal numbers are listed in Table 1. All female rats were monitored for phase of estrous cycle using vaginal cytology during the week prior to behavioral testing, 2-3 hours before behavioral testing, and on the day of sacrifice as described.^15,44^ All procedures were performed in accordance with the rules and regulations of the Institutional Animal Care and Use Committee of Drexel University College of Medicine and followed the NIH *Guide for the Care and Use of Animals*.

**Table 1.**
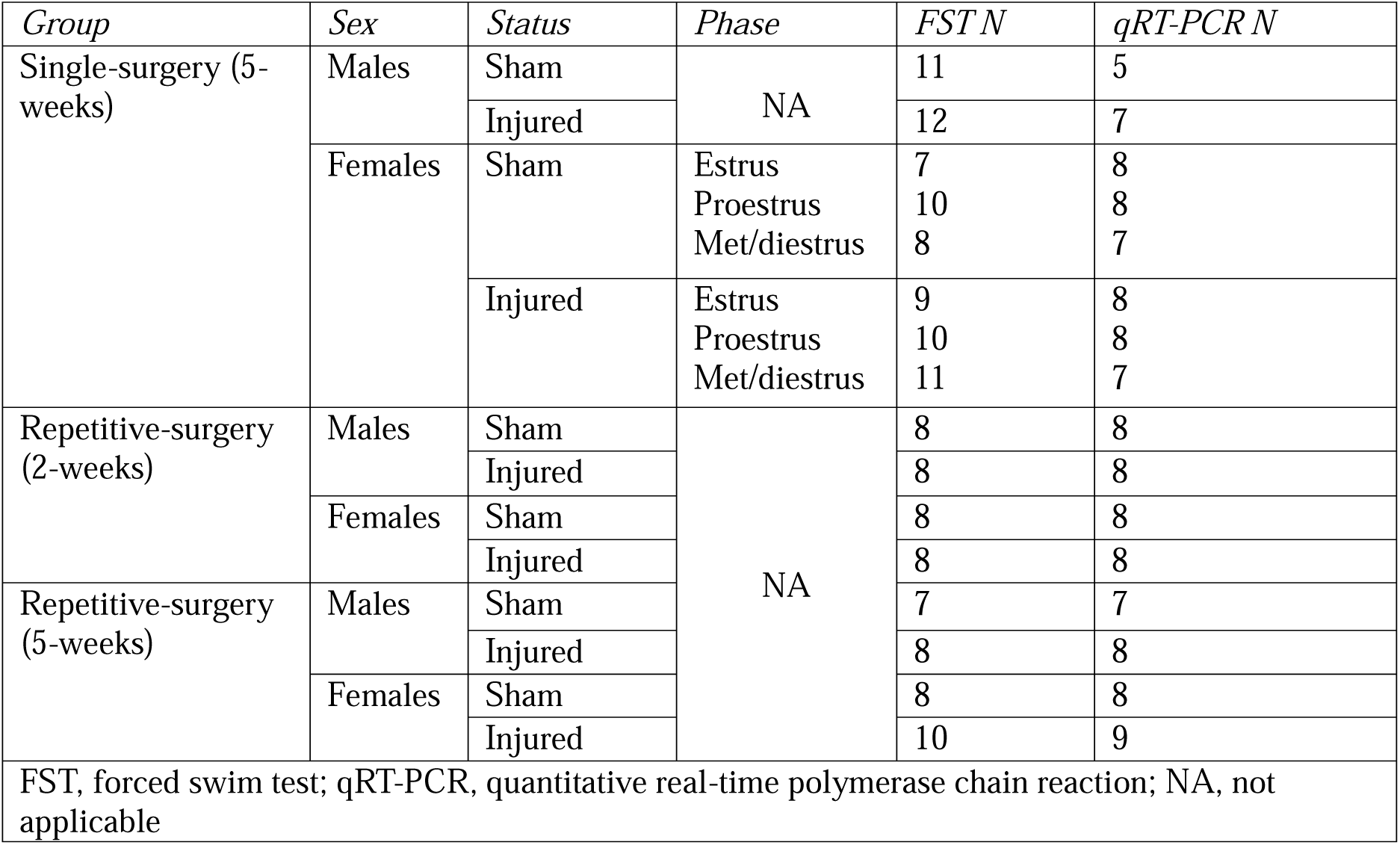
Number of rats used in each experimental group.

### Brain injury

Brain injuries were induced using an electronically driven controlled cortical impact device (Custom Design and Fabrication, Richmond, VA) as described.^15,16^ Impacts were induced using a convex metal indenter tip (5 mm diameter) which was fired to 2 mm below zero point (surface of the skull, 5.5 m/s velocity, 100 ms dwell time). Sham-injured animals were anesthetized but did not receive an impact. Immediately following impact, the rat was monitored for the time to regain their first breath (apnea) and time to regain normal posture when placed in the supine position (righting reflex). Starting at 6 weeks of age, single injured rats received only one sham- or brain-injury, whereas repetitive injured rats received three sham- or brain-injuries, each one spaced 3 days apart.

### Forced Swim Test (FST)

Sham- and brain-injured animals were tested for depression-like behavior using the FST,^45,46^ at 5 weeks after single injury and 2 or 5 weeks after repetitive injury.^15,16^ Rats swam for 10 continuous minutes in the apparatus (49 cm tall, 29 cm diameter acrylic cylinder filled with 30 cm of 25 ± 2°C water) as described.^47^ The time spent swimming (moving in the cylinder using hindlimbs), climbing (upward motion with forelimbs) and immobile (least amount of movement necessary to keep afloat) during the last 5 minutes of the test was quantified by an experimenter blind to injury status and estrous phase.

### Quantitative Real-Time PCR (qRT-PCR)

Rats were sacrificed one week following FST testing (single injury, based on estrous phase) or 2 days following FST (repetitive injury), during the first half of their light cycle. The NAc and the posterior PVT were dissected out to examine mRNA levels of CRF, DYN, ENK, (NAc) and PACAP (PVT), as described.^35,43^ Primer sequences and concentrations are listed in Table 2. Total RNA yield resulted in A_260_/A_280_ ratios between 1.90 and 2.20, indicating high purity. Target gene expression was quantified relative to cyclophilin-A using the relative quantification method (ΔΔC_T_).

**Table 2.**
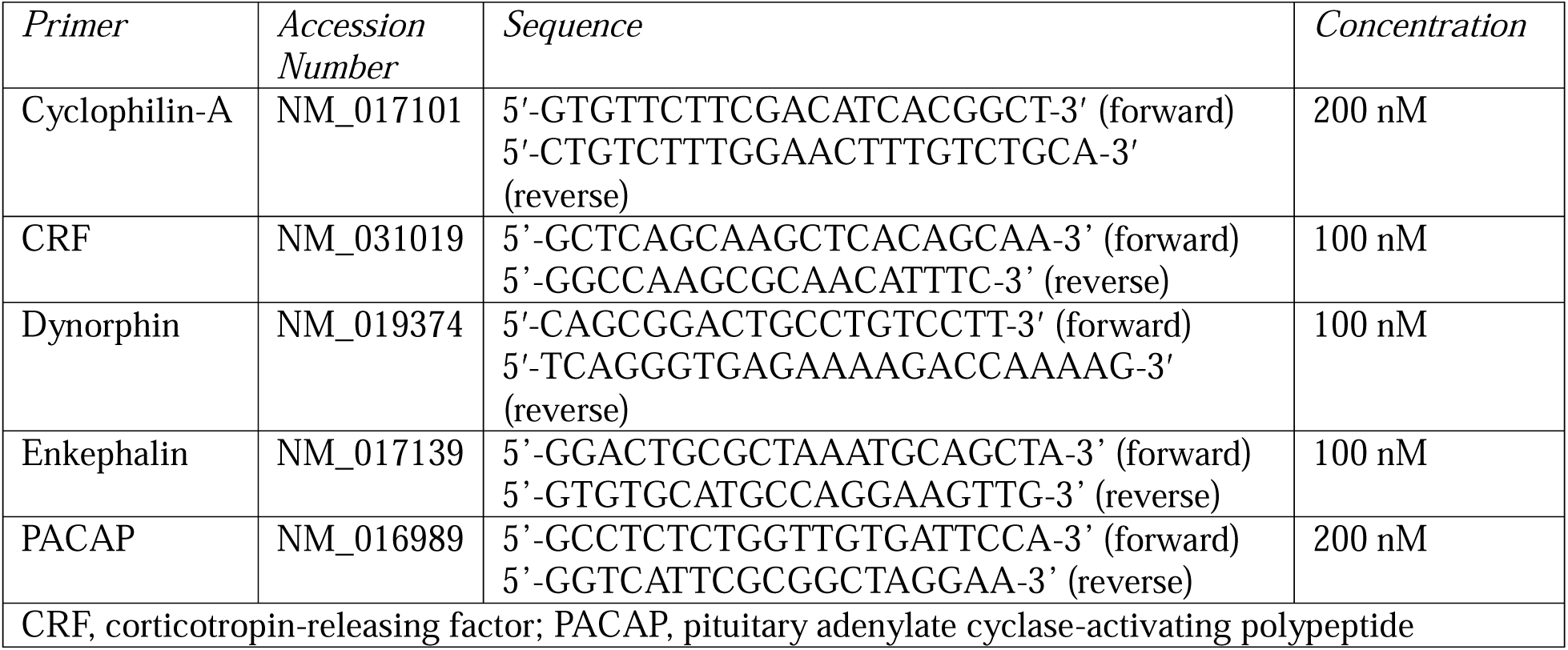
Primer sequences and concentrations used for quantitative real-time polymerase chain reaction experiments.

### Statistics

Statistical tests were performed using Statistica 7 (StatSoft, Tulsa OK). Body weights, latencies to regain righting reflex and apnea times during surgeries were compared using a two-way analysis of variance (ANOVA) with sex and injury status as independent variables. For repetitive surgeries, repeated measures ANOVA were used to compare these variables between each surgery day. Chi-Square tests were used to analyze fractures and hematomas within the brain-injured rats acutely following injury. FST behavior (time spent swimming, climbing and immobile) and qRT-PCR (ΔΔC_T_ values) data were compared using a 2-way ANOVA with sex and injury status as independent variables. For females subjected to single surgeries, behavioral and qRT-PCR data were further analyzed using a 2-way ANOVA with injury status and estrous phase as independent variables, except for comparing PACAP expression in females during estrus, where a *t*-test was used. *Post-hoc* analysis using the Newman-Keuls test were performed when significant effects were found (*p* ≤ 0.05). As an *a priori* assumption was made that sex-related differences in PACAP expression could exist in females,^35^ Newman-Keuls comparison tests were used to examine injury effects within each sex, even when interaction effects between sex and injury status did not reach significance. Behavioral and qRT-PCR data are presented as percentage of sham values. All data are presented as mean ± standard error of the mean (SEM).

## Results

### Acute responses to injury

A single impact to the closed skull of 6-week-old Long-Evans rats resulted in increased latency to recover righting reflexes compared to sham animals (*F*_1,87_ = 144.22, *p* = 0.000001, Table 3). Brain-injured animals experienced apnea (1-13 s) immediately after injury (*F*_1,87_ = 209.52, *p* = 0.000001, Table 3). Males displayed a greater loss of righting reflex (*p* = 0.0002) and apnea time (*p* = 0.001) compared to females. For male and female rats, impact was associated with a moderate or mild hematoma (91%) and a small linear fracture on the skull (98%).

**Table 3.**
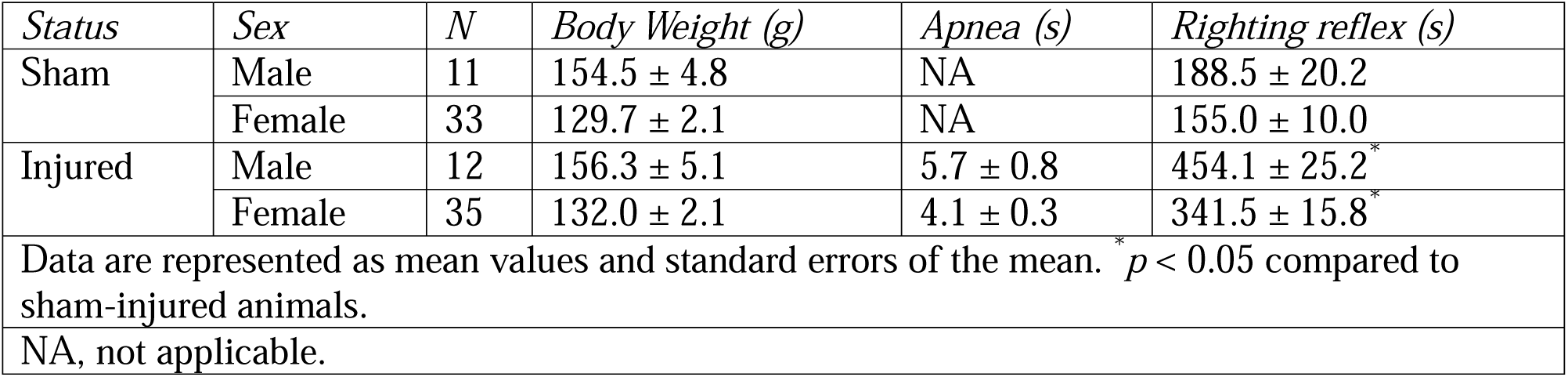
Acute neurological responses to a single impact on the intact skull of adolescent male and female Long-Evans rats.

Repetitive impacts also resulted in increased latency to recover righting reflexes compared to sham animals (*F*_1,71_ = 54.82, *p* = 0.000001, Table 4). With a significant interaction between injury status and surgery day (*F*_2,142_ = 5.43, *p* = 0.005), the loss of righting reflex was greater for brain-injured animals on the first day of injury compared to injury days 2 (*p* = 0.0005) and 3 (*p* = 0.0003); no differences in loss of righting reflex were observed in brain-injured animals between days 2 and 3 or in sham-injured animals across all surgery days. Brain-injured animals experienced apnea (1-7 s) immediately following injuries (*F*_1,71_ = 349.62, *p* = 0.000001, Table 4). There were no differences in apnea times between the three injury days. For male and female rats, impact was associated with a moderate or mild hematoma (98%) and a small linear fracture after the first injury (76%).

**Table 4.**
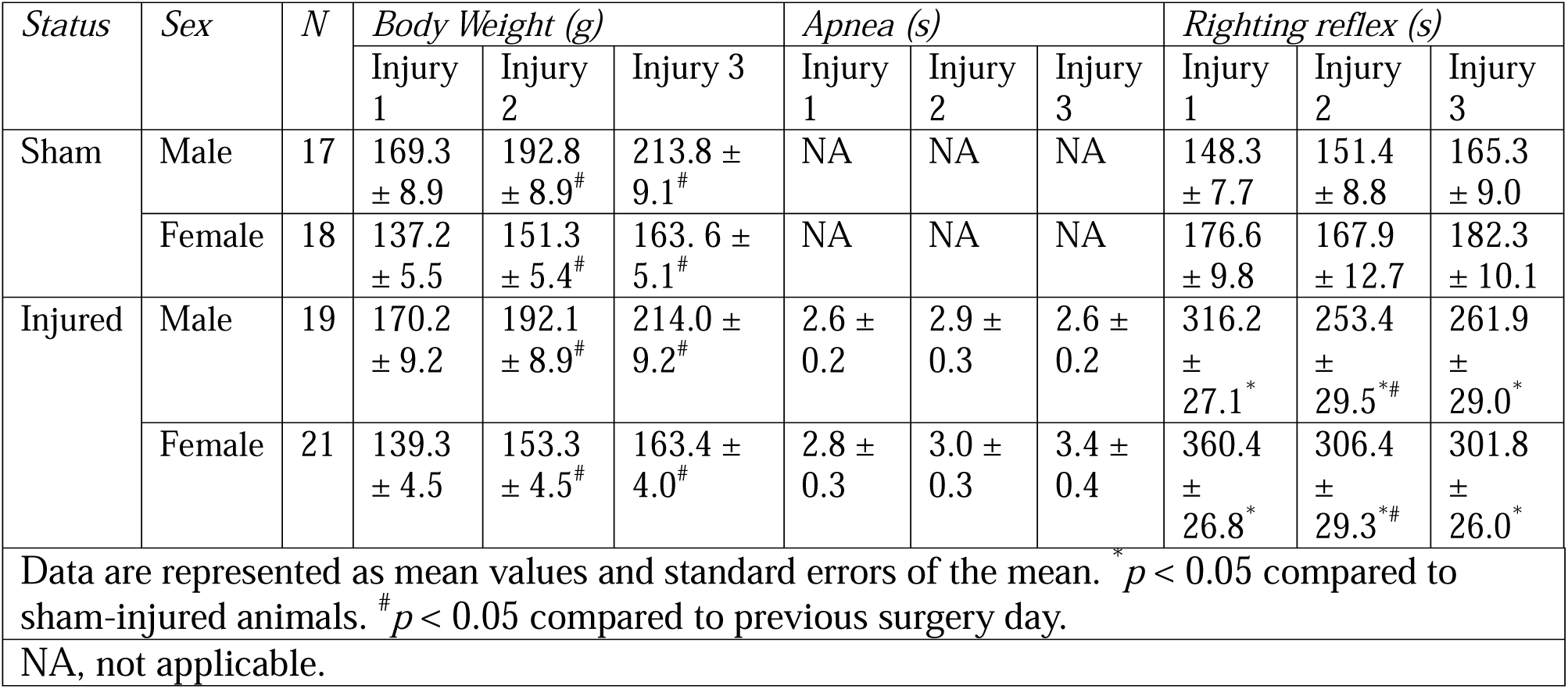
Acute neurological responses to repetitive impact on the intact skull of adolescent male and female Long-Evans rats.

Sham- and brain-injured rats displayed irregular estrous cycles for 2 weeks after surgery or repetitive mTBI, respectively, which returned to the normal 4–5-day cycle by 5 weeks.

### Depression-like behavior in the FST

When tested in the FST at 5 weeks following a single mTBI, there was no effect of injury status on time spent immobile (Figure 1A), swimming (Figure 1B), or climbing (Figure 1C). However, there was a significant sex effect, with females spending more time immobile (*F*_1,70_ = 44.27, *p* = 0.0001) and less time swimming than males (*F*_1,67_ = 56.60, *p* = 0.0001). Evaluating females in different estrous phases revealed a significant effect of injury status (*F*_1,44_ = 7.01, *p* = 0.01) and an injury status and estrous phase interaction (*F*_2,44_ = 8.06, *p* = 0.001), but no effect of estrous phase (*F*_2,44_ = 1.89, *p* = 0.16) on time spent immobile (Figure 1D). *Post-hoc* analysis revealed that brain-injured females spent more time immobile than their sham-injured counterparts in the estrus phase (*p* = 0.0003) and that sham-injured rats in met/diestrus (*p* = 0.02) and proestrus (*p* = 0.005) spent more time immobile than sham-females in estrus. For time spent swimming, injury status and estrous phase interaction neared significance, where injured females during estrus spent less time swimming (*F*_2,44_ = 3.17, *p* = 0.052, Figure 1E). Females during estrus spent more time climbing than females in other phases (*F*_2,43_ = 6.12, *p* = 0.005, Figure 1F).

**Figure 1.**
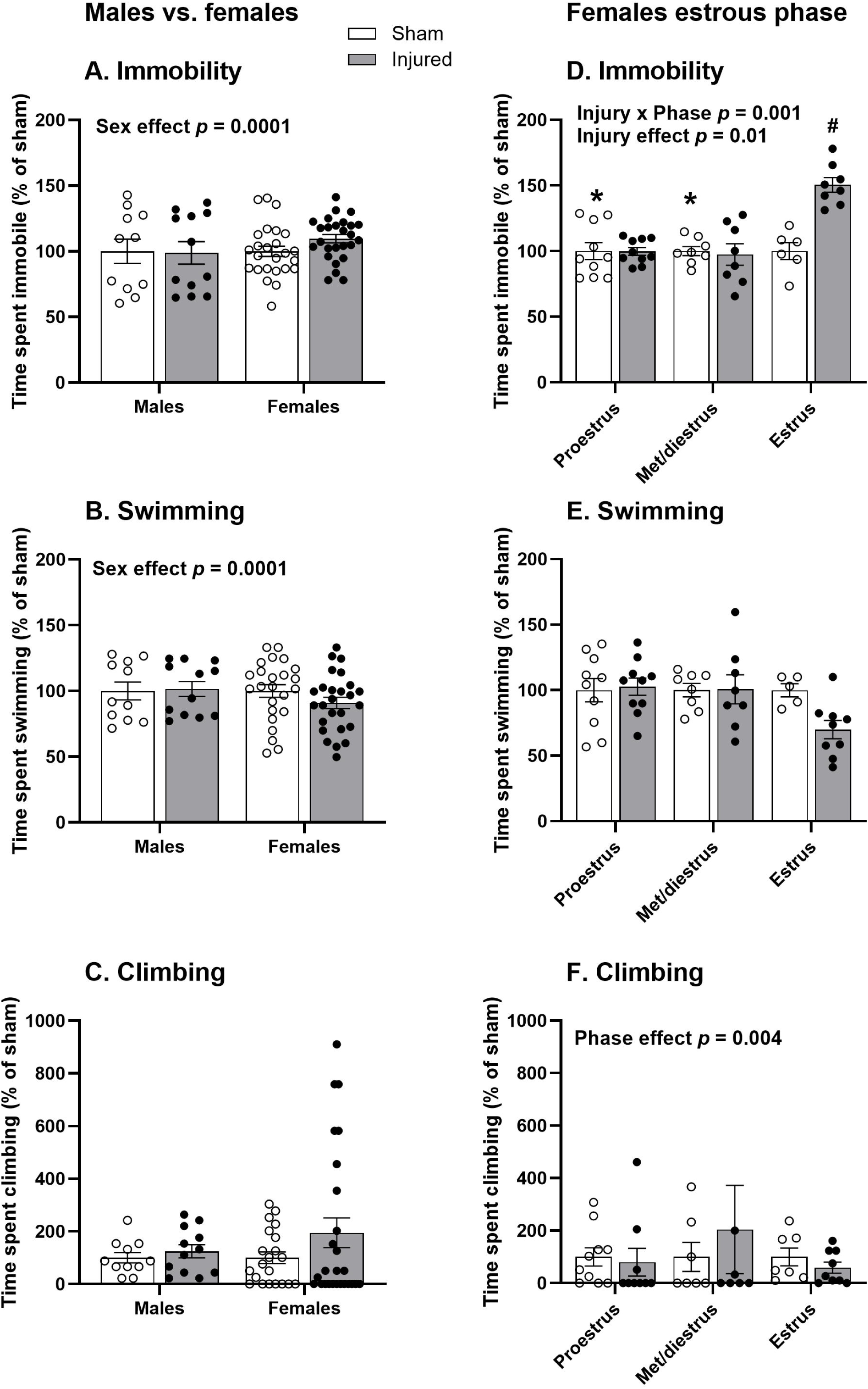
FST behavior 5 weeks following a single mTBI or sham-injury in male and female Long Evans rats **(A-C)** and females separated by estrous phase at the time of testing **(D-F)**. Scatterplots of immobility **(A,D)**, swimming **(B,E)** and climbing **(C,F)** times of sham-(open circles) and brain-injured (closed circles) rats. Bars represent mean percent of sham value for each group and standard error of the mean. *,# *p* < 0.05 compared to sham estrus. Statistical analyses were performed on time spent in the various behaviors, but data are plotted as a function of sham values.

At 2 weeks following repetitive mTBI, injury status had no significant effect on immobility (Figure 2A), swimming (Figure 2B) or climbing (Figure 2C). Females spent more time immobile (*F*_1,27_ = 4.62, *p* = 0.04) and less time swimming (*F*_1,28_ = 9.02, *p* = 0.006) than males overall, with no significant difference in time spent climbing. At 5 weeks following repetitive mTBI, there was a significant effect of injury status (*F*_1,28_ = 5.57, *p* = 0.02) and *post-hoc* analysis of an injury status and sex interaction on time spent immobile (*F*_1,28_ = 5.38, *p* = 0.03, Figure 2D) revealed that brain-injured females spent more time immobile than sham females (*p* = 0.01) and injured males (*p* = 0.01). Brain-injured rats overall spent significantly less time swimming than sham-injured (*F*_1,28_ = 14.55, *p* = 0.0007, Figure 2E) and females spent less time climbing than males (*F*_1,27_ = 4.67, *p* = 0.04, Figure 2F).

**Figure 2.**
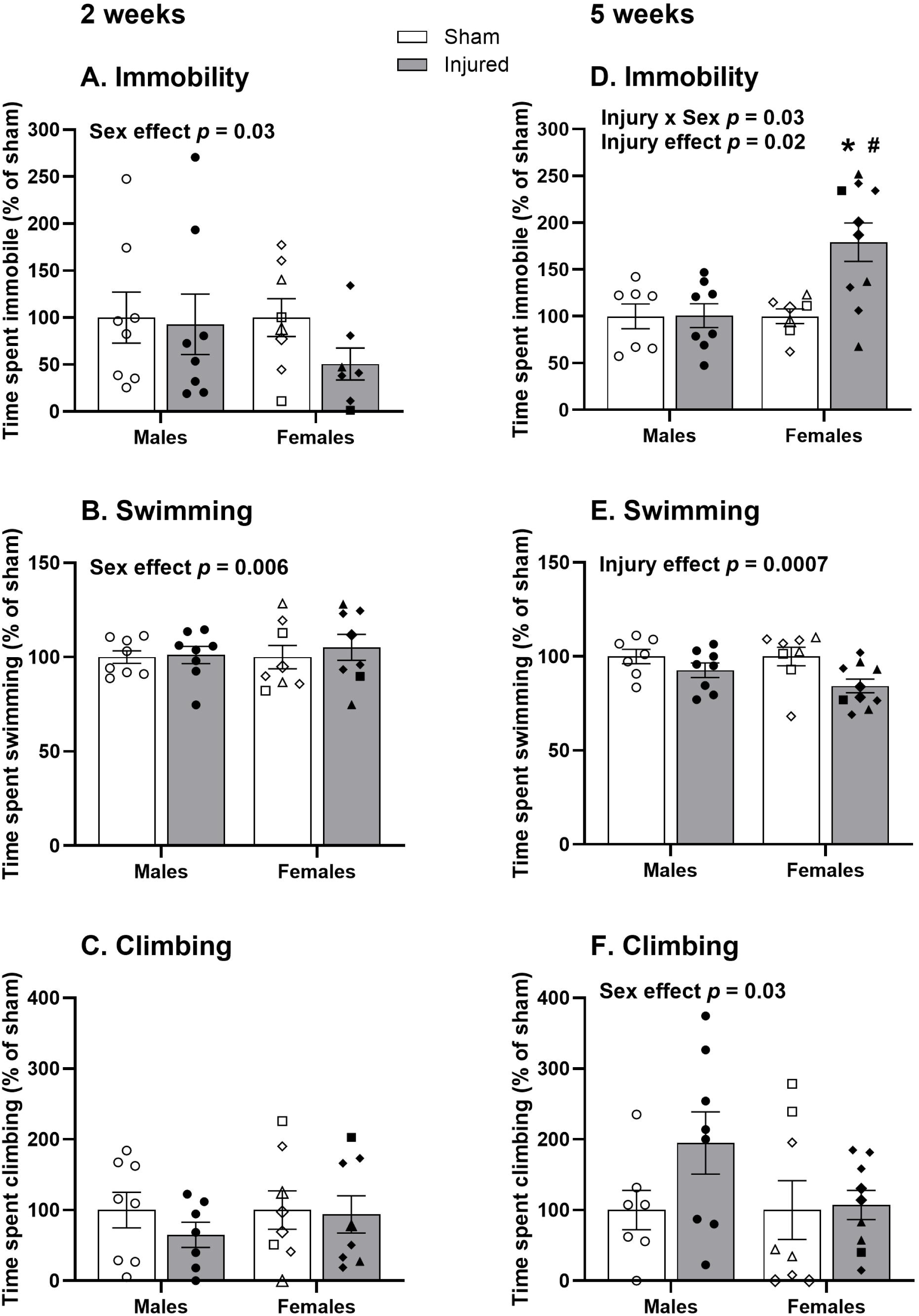
FST behavior in male and female Long-Evans rats at 2 weeks **(A-C)** and 5 weeks **(D-F)** following repetitive mTBI or sham surgery. Scatterplots of immobility **(A,D)**, swimming **(B,E)** and climbing **(C,F)** times of sham-(open symbols) and brain-injured (closed symbols) rats. Bars represent mean percent of sham value for each group and standard error of the mean. Symbol key: square estrus phase; triangle proestrus phase; diamond met/diestrus phases. *, p < 0.05 compared to sham females. # *p* < 0.05 compared to injured males. Statistical analyses were performed on time spent in the various behaviors, but data are plotted as a function of sham values.

### Neuropeptide mRNA in the NAc

Five weeks following a single mTBI, female rats displayed higher expression of CRF compared to males (*F*_1,53_ = 4.84, *p* = 0.03) but there was no change in CRF as an effect of injury (Figure 3A). Additionally, DYN (*F*_1,53_ = 7.91, *p* = 0.007, Figure 3B) and ENK (*F*_1,53_ = 37.92, *p* = 0.0001, Figure 3C) expression was decreased in females compared to males. When comparing females by estrous phase, a significant effect of injury status (*F*_1,40_ = 4.03, *p* = 0.05, Figure 3D) showed that injured females had reduced CRF expression compared to shams. A *post-hoc* analysis of estrous phase (*F*_2,40_ = 5.91, *p* = 0.006) revealed that females in estrus (*p* = 0.006) and met/diestrus phases (*p* = 0.03) had significantly reduced CRF mRNA expression compared to proestrus. There was a significant decrease in DYN mRNA expression in injured females compared to shams regardless of estrous phase (*F*_1,40_ = 4.15, *p* = 0.04, Figure 3E). There were no differences in ENK mRNA expression (Figure 3F).

**Figure 3.**
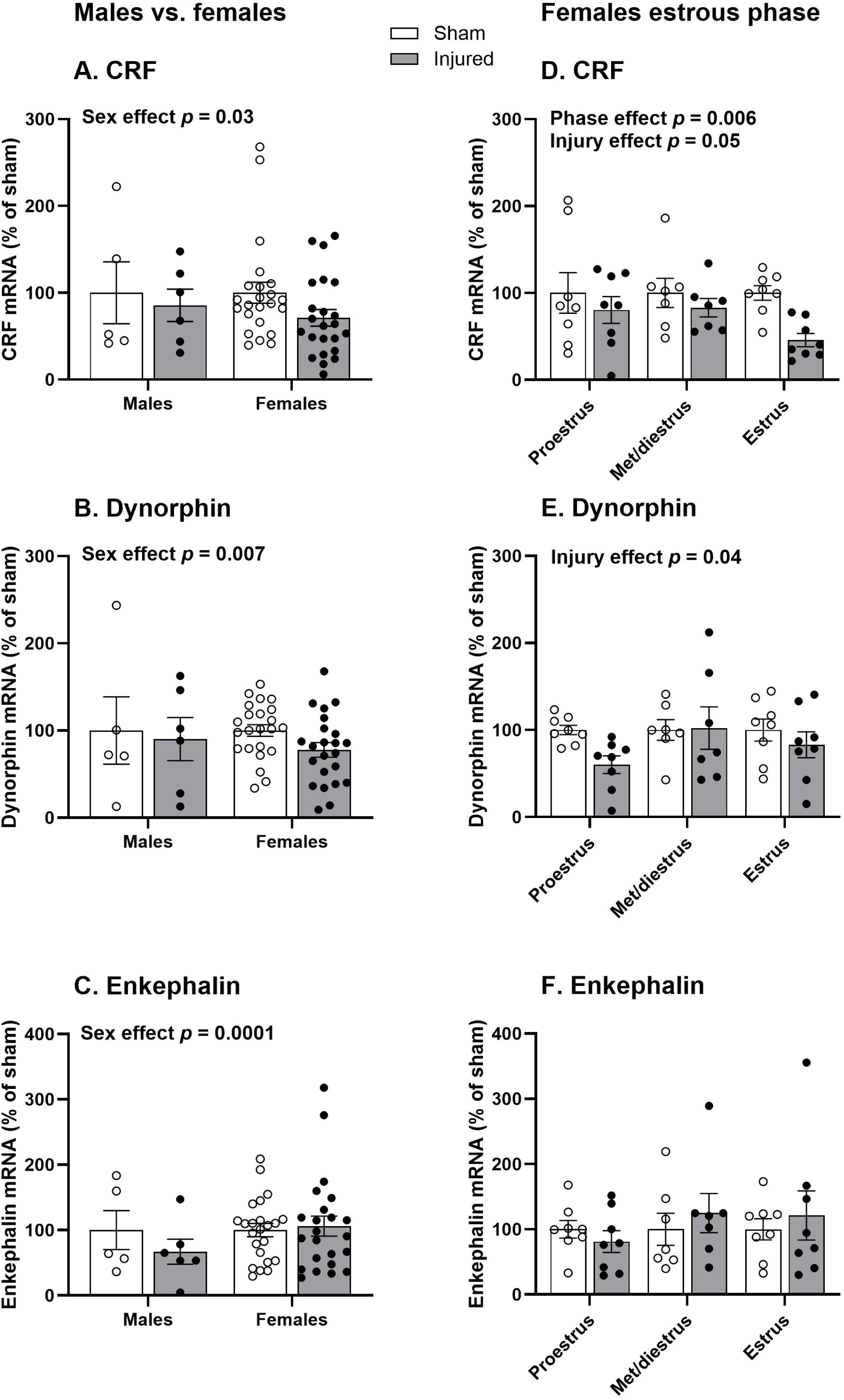
Neuropeptide mRNA expression in the nucleus accumbens (NAc) 5 weeks after a single mTBI or sham surgery in male and female Long-Evans rats **(A-C)** and in females separated by estrous phase at the time of tissue collection **(D-F)**. Scatterplots of corticotropin-releasing factor (CRF) **(A,D)**, dynorphin **(B,E)** and enkephalin **(C,F)** mRNA expression of sham-(open circles) and brain-injured (closed circles) rats. Bars represent mean percent of sham value for each group and standard error of the mean. Statistical analyses were performed on ΔΔC_T_ values, but data are plotted as a function of sham values.

Five weeks after repetitive mTBI, there were no significant differences regardless of injury status or sex in CRF expression in the NAc (Figure 4A). In contrast, females showed less DYN mRNA expression in the NAc compared to males (*F*_1,26_ = 10.91, *p* = 0.003, Figure 4B) and brain-injured rats overall showed a significant decrease of ENK expression compared to sham-injured rats (*F*_1,26_ = 4.30, *p* = 0.05, Figure 4C).

**Figure 4.**
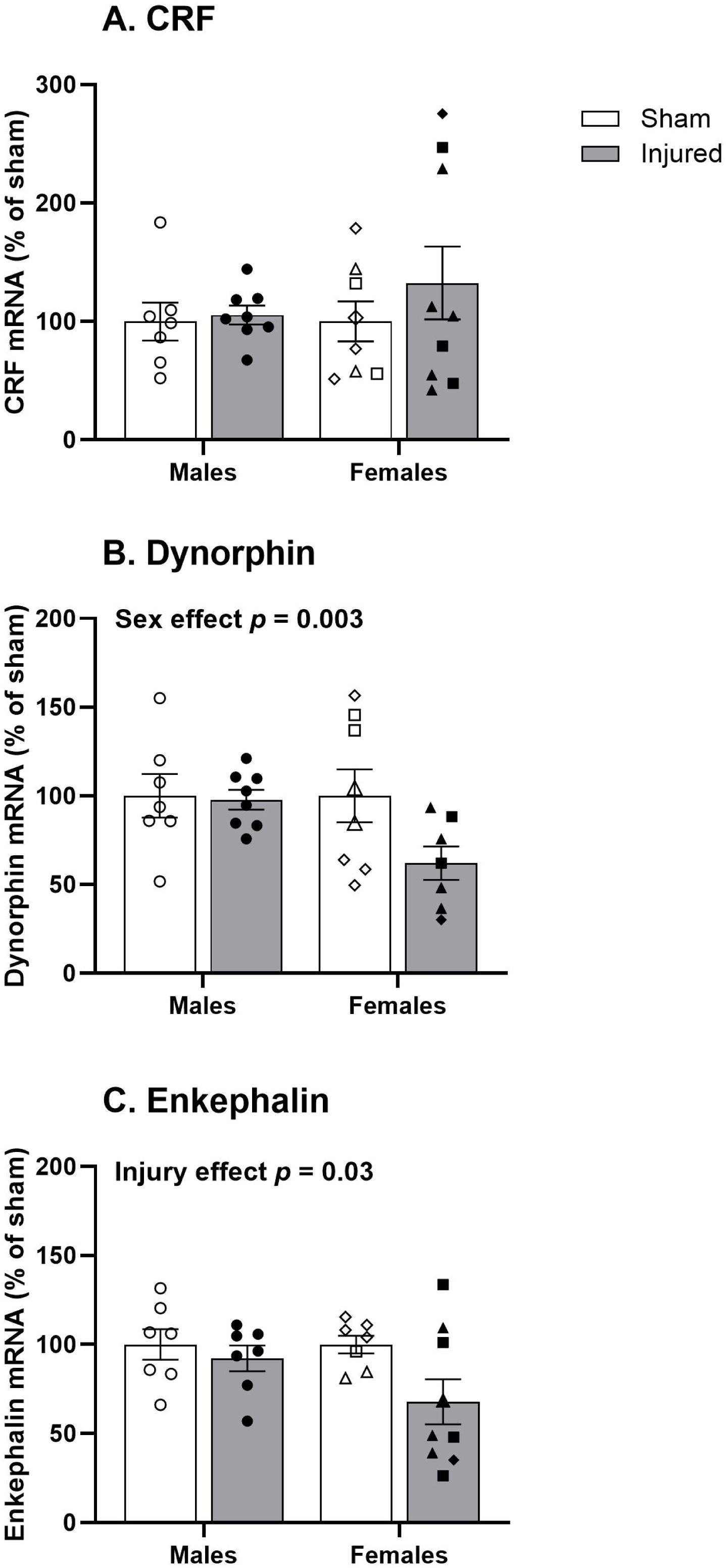
Neuropeptide mRNA expression in the nucleus accumbens 5 weeks after repetitive mTBI or sham surgery in male and female Long-Evans rats. Scatterplots of corticotropin-releasing factor (CRF) **(A)**, dynorphin **(B)** and enkephalin **(C)** mRNA expression of sham-(open symbols) and brain-injured (closed symbols) rats. Bars represent mean percent of sham value for each group and standard error of the mean. Symbol key: square estrus phase; triangle proestrus phase; diamond met/diestrus phases. Statistical analyses were performed on ΔΔC_T_ values, but data are plotted as a function of sham values.

### PACAP mRNA in the PVT

Five weeks after a single mTBI, brain-injured females during estrus did not display a difference in PACAP mRNA expression compared to shams (Figure 5A). Two weeks after repetitive mTBI, females compared to males showed greater expression of PACAP mRNA (*F*_1,27_ = 9.55, *p* = 0.005). *Post-hoc* analysis of a significant interaction between injury status and sex (*F*_1,27_ = 6.88, *p* = 0.01, Figure 5B) revealed that injured females had significantly greater expression of PVT PACAP mRNA compared to sham females (*p* = 0.004) and injured males (*p* = 0.002). Five weeks after repetitive injury, injured rats displayed higher expression of PACAP mRNA regardless of sex (*F*_1,26_ = 6.22, *p* = 0.02, Figure 5C). However, injured females showed a higher increase in PACAP (+163%, *η^2^_p_* = 0.25) than males (+29%, *η^2^_p_* = 0.02) compared to shams, and *post-hoc* comparisons revealed that higher expression in injured rats was driven by a significant difference in sham/injured females (*p* = 0.04).

**Figure 5.**
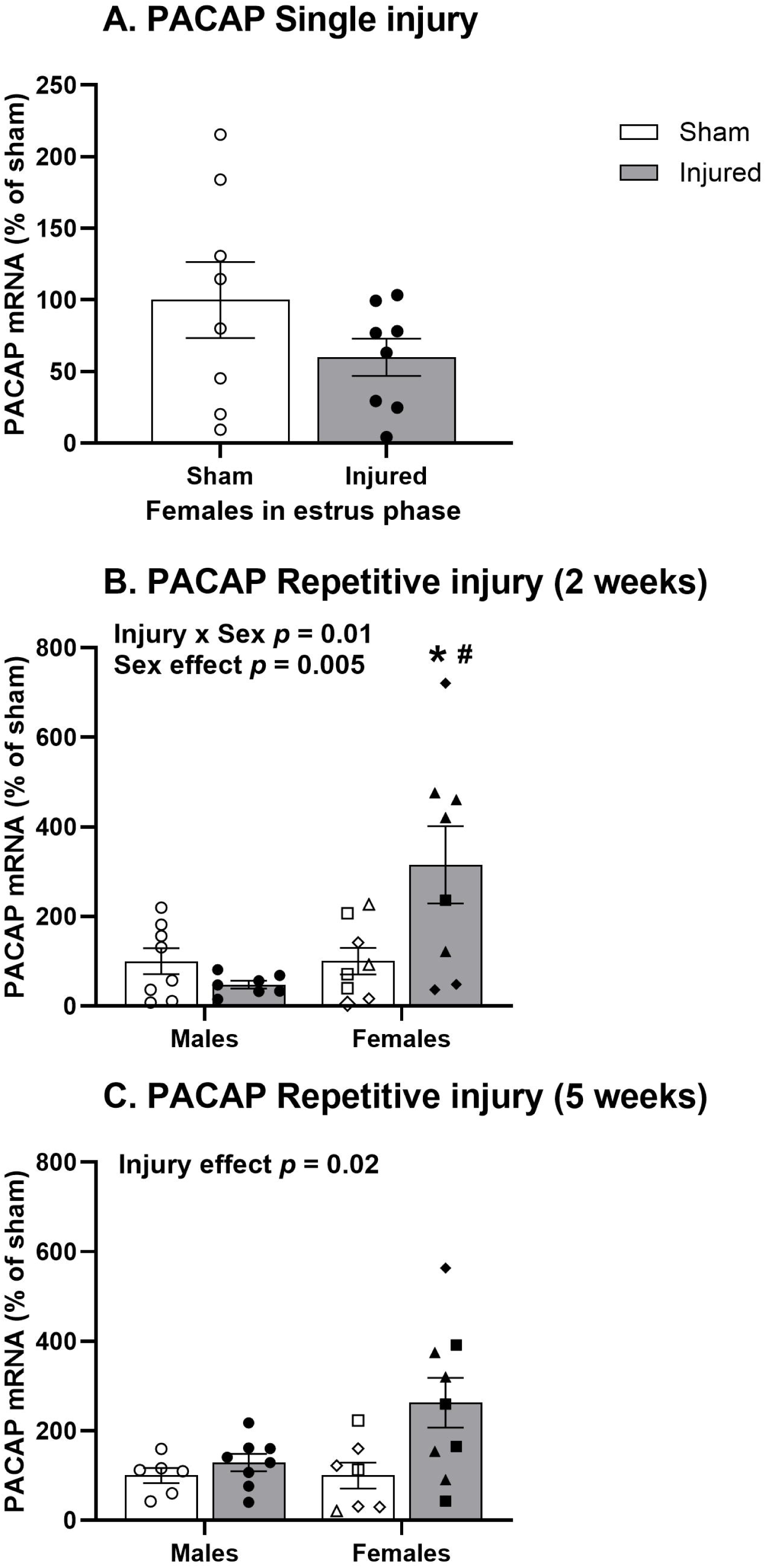
Pituitary adenylate cyclase-activating polypeptide (PACAP) mRNA expression in the paraventricular nucleus of the thalamus 5 weeks after a single surgery in female rats during estrus phase at the time of tissue collection **(A)** or 2 weeks **(B)** and 5 weeks **(C)** after repetitive mTBI or sham surgery in male and female Long-Evans rats. Scatterplots of PACAP mRNA expression of sham-(open symbols) and brain-injured (closed symbols) rats. Bars represent mean percent of sham value for each group and standard error of the mean. Symbol key: square estrus phase; triangle proestrus phase; diamond met/diestrus phases. * *p* < 0.05 compared to sham females. # *p* < 0.05 compared to injured males. Statistical analyses were performed on ΔΔC_T_ values, but data are plotted as a function of sham values.

## Discussion

In this study, we found that female Long-Evans rats showed increased depression-like behavior in the FST after single and repetitive mTBI. In single-but not repetitive-injured animals, this increase was dependent on estrus phase. Following single injury, CRF and DYN mRNA expression in the NAc was reduced in injured females with the decrease in CRF most pronounced during the estrus phase. In contrast, there were no significant changes in CRF or DYN expression following repetitive injury. Within the PVT, PACAP expression was only increased in females following repetitive mTBI. These differential changes underscore the importance of sex, estrous cycle and injury severity in governing changes in behavior and associated molecular responses.

The present data in Long-Evans rats recapitulates our previous work in Sprague-Dawley rats,^15,16^ suggesting that strain differences may not be prevalent following mTBI. It has been reported that female rats in the estrus phase display increased immobility even after treatment with anti-depressants.^48,49^ Similarly, women may experience worse depression symptoms and higher rates of suicide when estrogen levels are lowest.^50^ However, it appears that phases of the estrous cycle may not govern FST behavior following repeated mTBI.^16^ We observed that repetitive mTBI increased immobility time only at the 5 week timepoint post-injury and not at the 2-week time point, similar to what was observed following single injury.^15^ Previous studies in rats injured at 30 days of age have reported significant differences earlier than 5 weeks post-injury,^17,51^ which may be explained as a function of age-at-injury (early versus late adolescence).^52^ In this regard, repetitive mTBI in 48 day old mice (late adolescence) did not result in depression-like behavior.^18^

Five weeks after single mTBI, injured females showed reduced mRNA expression of CRF and DYN in the NAc. Estrous phase did not significantly affect DYN expression but a decrease in CRF expression was the most striking among injured females in estrus, when females also spent more time immobile in the FST. These observations suggest that CRF may interact with both female ovarian hormones and injury, while DYN expression may only change in females as a function of injury, independent of estrous cycling. Previous work has identified increased grooming behavior in females treated with CRF during proestrus, suggesting an interaction between CRF and higher levels of ovarian hormones.^41^ This may explain why single-injured females during their estrus phase display increased immobility and decreased CRF: ovarian hormones interact with CRF to produce active behaviors in female rats, and injury may cause a change in CRF synthesis that dysregulates the natural balance. Interestingly, neither DYN nor CRF expression was altered following repetitive TBI. However, ENK mRNA in the NAc and PACAP mRNA in the PVT only changed following repetitive injury. Other studies have observed a downregulation of ENK in the NAc that accompanied increased anhedonia or susceptibility to stress in rats and mice.^32,33^ PACAP overexpression in the PVT of female rats reduced time spent swimming in the FST.^47^ Although we observed PACAP upregulation in the PVT after both 2 and 5 weeks, this suggests that it may be causal of the observed FST behavior 5 weeks after repetitive injury, as the behavioral effects of PACAP may be age-dependent.^53^ These opposing limbic neuropeptide mRNA changes after single or repetitive mTBI highlight possible mechanistic differences in mTBI-induced regulation of behavioral changes specific to females.

A limitation of this study is that only one behavior test (FST) was employed to observe depression-like behavior. Although immobility behavior is commonly referred to as “depression-like” behavior, recent commentaries suggest that FST may instead evaluate coping behavior.^54,55^ Other studies report that mTBI induces affective behavioral deficits, such as anxiety-like behavior and anhedonia,^17,56–58^ we have not observed anxiety-like behaviors following either single or repetitive injuries in adolescent female rats.^15,16^ This study also only begins to look at chronic changes in the expression of a few different limbic region neuropeptides. Some studies have focused on the therapeutic potential of treating emotional dysregulation following brain injury with CRF and DYN receptor antagonists.^23,25^ Further research is needed into affective behavioral deficits and the mechanisms of these deficits following adolescent mTBI, with more consideration of females and the repetitive nature of sport-related concussions.

### Conclusion

Our study shows that both single and repetitive mild midline impacts during adolescence result in chronic depression-like behavior that is specific to female Long-Evans rats and that this is only dependent on the estrous cycle following a single injury. It also reveals differences in level of expression of different limbic neuropeptides after single or repetitive impacts in females, similarly only estrous cycle dependent after single injury. These observations suggest that chronic depression-like behavior and limbic neuropeptide changes following adolescent mTBI may be specific to females and only dependent on estrous cycling after single injury. There is a need to develop more appropriate treatment for TBI-induced depression that takes these sex-specific and injury-type differences into consideration.

## Acknowledgements

The authors thank Lilia Sanzalone for her assistance with qRT-PCR analyses and Dana Lengel and Moriah Harton for their assistance with injury surgeries and behavioral assessments.

